# The SARS-CoV-2 receptor, Angiotensin converting enzyme 2 (ACE2) is required for human endometrial stromal cell decidualization

**DOI:** 10.1101/2020.06.23.168252

**Authors:** Sangappa B. Chadchan, Vineet K. Maurya, Pooja Popli, Ramakrishna Kommagani

**Affiliations:** Center for Reproductive Health Sciences, Department of Obstetrics and Gynecology Washington University School of Medicine, St. Louis, MO, 63110, USA

**Keywords:** SARS-CoV-2, Endometrium, Embryo implantation, Stromal cells, Decidualization

## Abstract

**STUDY QUESTION:** Is SARS-CoV-2 receptor, angiotensin-converting enzyme 2 (ACE 2) expressed in the human endometrium during the menstrual cycle, and does it participate in endometrial decidualization?

**SUMMARY ANSWER:** ACE2 protein is highly expressed in human endometrial stromal cells during the secretory phase and is essential for human endometrial stromal cell decidualization.

**WHAT IS KNOWN ALREADY:** ACE2 is expressed in numerous human tissues including the lungs, heart, intestine, kidneys and placenta. ACE2 is also the receptor by which SARS-CoV-2 enters human cells.

**STUDY DESIGN, SIZE, DURATION:** Proliferative (n = 9) and secretory (n = 6) phase endometrium biopsies from healthy reproductive-age women and primary human endometrial stromal cells from proliferative phase endometrium were used in the study.

**PARTICIPANTS/MATERIALS, SETTING, METHODS:** ACE2 expression and localization were examined by qRT-PCR, Western blot, and immunofluorescence in both human endometrial samples and mouse uterine tissue. The effect of *ACE2* knockdown on morphological and molecular changes of human endometrial stromal cell decidualization were assessed. Ovariectomized mice were treated with estrogen or progesterone to determine the effects of these hormones on ACE2 expression.

**MAIN RESULTS AND THE ROLE OF CHANCE:** In human tissue, ACE2 protein is expressed in both endometrial epithelial and stromal cells in the proliferative phase of the menstrual cycle, and expression increases in stromal cells in the secretory phase. The *ACE2* mRNA (*P* < 0.0001) and protein abundance increased during primary human endometrial stromal cell (HESC) decidualization. HESCs transfected with *ACE2*-targeting siRNA were less able to decidualize than controls, as evidenced by a lack of morphology change and lower expression of the decidualization markers *PRL* and *IGFBP1* (*P* < 0.05). In mice during pregnancy, ACE2 protein was expressed in uterine epithelial and stromal cells increased through day six of pregnancy. Finally, progesterone induced expression of *Ace2* mRNA in mouse uteri more than vehicle or estrogen (*P* < 0.05).

**LARGE SCALE DATA:** N/A.

**LIMITATIONS, REASONS FOR CAUTION:** Experiments assessing the function of ACE2 in human endometrial stromal cell decidualization were *in vitro*. Whether SARS-CoV-2 can enter human endometrial stromal cells and affect decidualization have not been assessed.

**WIDER IMPLICATIONS OF THE FINDINGS:** Expression of ACE2 in the endometrium allow SARS-CoV-2 to enter endometrial epithelial and stromal cells, which could impair *in vivo* decidualization, embryo implantation, and placentation. If so, women with COVID-19 may be at increased risk of early pregnancy loss.

**STUDY FUNDINGS/COMPETING INTEREST(S):** This study was supported by National Institutes of Health / National Institute of Child Health and Human Development grants R01HD065435 and R00HD080742 to RK and Washington University School of Medicine start-up funds to RK. The authors declare that they have no conflicts of interest.

## Introduction

Although much of the focus during the severe acute respiratory syndrome coronavirus 2 (SARS-CoV-2)/coronavirus disease 2019 (COVID-19) pandemic has been on respiratory symptoms, some reports suggest that SARS-CoV-2 and the related Middle East Respiratory Syndrome Coronavirus can cause pregnancy complications such as pre-term birth and miscarriages (Favre et al. 2020). Additionally, a few reports noted that pregnant women with COVID-19 had maternal vascular malperfusion and decidual arteriopathy in their placentas (Schwartz and Dhaliwal 2020; Shanes et al. 2020a), and a recent clinical case study reported a second-trimester miscarriage in a woman with COVID-19 (Baud et al. 2020). However, whether SARS-CoV-2 infects the uterus has not been determined.

It seems likely that SARS-CoV-2 could infect the uterus because its receptor, Angiotensin Converting Enzyme 2 (ACE2), is expressed fairly ubiquitously in human tissues such as the lungs, heart, intestine, kidneys, and placenta (Hamming et al. 2004; Harmer et al. 2002; Riviere et al. 2005). Moreover, ACE2 functions by cleaving the vasoconstrictor angiotensin II to the vasodilator angiotensin (1-7). As a component of the renin–angiotensin system, ACE2 plays an important role in regulating maternal blood pressure during pregnancy. ACE2 is expressed in the rat uterus during mid- and late pregnancy (Merrill et al. 2002; Neves et al. 2008). In addition, ACE2 mRNA expression was noted in the uterus of both rats (Brosnihan et al. 2012) and humans (Vaz-Silva et al. 2009), in which its expression may be higher in the secretory phase than in the proliferative phase of the menstrual cycle (Vaz-Silva et al. 2009).

During the secretory phase, the uterine stromal cells prepare for embryo implantation by undergoing a progesterone-mediated differentiation process called decidualization. In this process, the stromal cells divide, change from a fibroblastic to an epithelioid morphology, and change their pattern of gene expression. Decidualization is essential for trophoblast invasion and placentation (Carson et al. 2000; Norwitz et al. 2001; Wilcox et al. 1999), and defects in this process may underlie early pregnancy loss in some women. Given the important function of the uterine stroma and the possibility that SARS-CoV-2 could infect the uterus, our goal here was to determine whether ACE2 is expressed in endometrial stromal cells, is regulated by progesterone, and is required for decidualization.

## Results and Discussion

We first sought to determine whether ACE2 is expressed in the endometrium and whether its expression differs according to the phase of the menstrual cycle. Thus, we obtained endometrial biopsies from women during the proliferative or secretory phase of the menstrual cycles and performed immunofluorescence with an ACE2-specific antibody. In the proliferative phase, ACE2 was highly expressed in epithelial cells than in stromal cells (**Fig. 1A**). However, in the secretory phase, ACE2 expression was increased in the stromal cells (**Fig. 1A**). Thus, we wondered whether ACE2 expression increased during *in vitro* decidualization of human endometrial stromal cells (HESCs). We isolated primary HESCs, exposed them to decidualizing conditions, and confirmed that expression of the decidualization markers Prolactin *(PRL)* and Insulin-like growth factor-binding protein-1 *(IGFBP1*) increased over six days. *ACE2* mRNA also increased over this time (**Fig. 2A**). Consistent with this finding, ACE2 protein abundance increased during decidualization, as shown by both immunoblotting (**Fig. 2B**) and immunofluorescence (**Fig. 2C**). As expected, ACE2 protein predominantly localized in the cytoplasm and cell membrane of decidualized HESCs.

**Figure 1:**
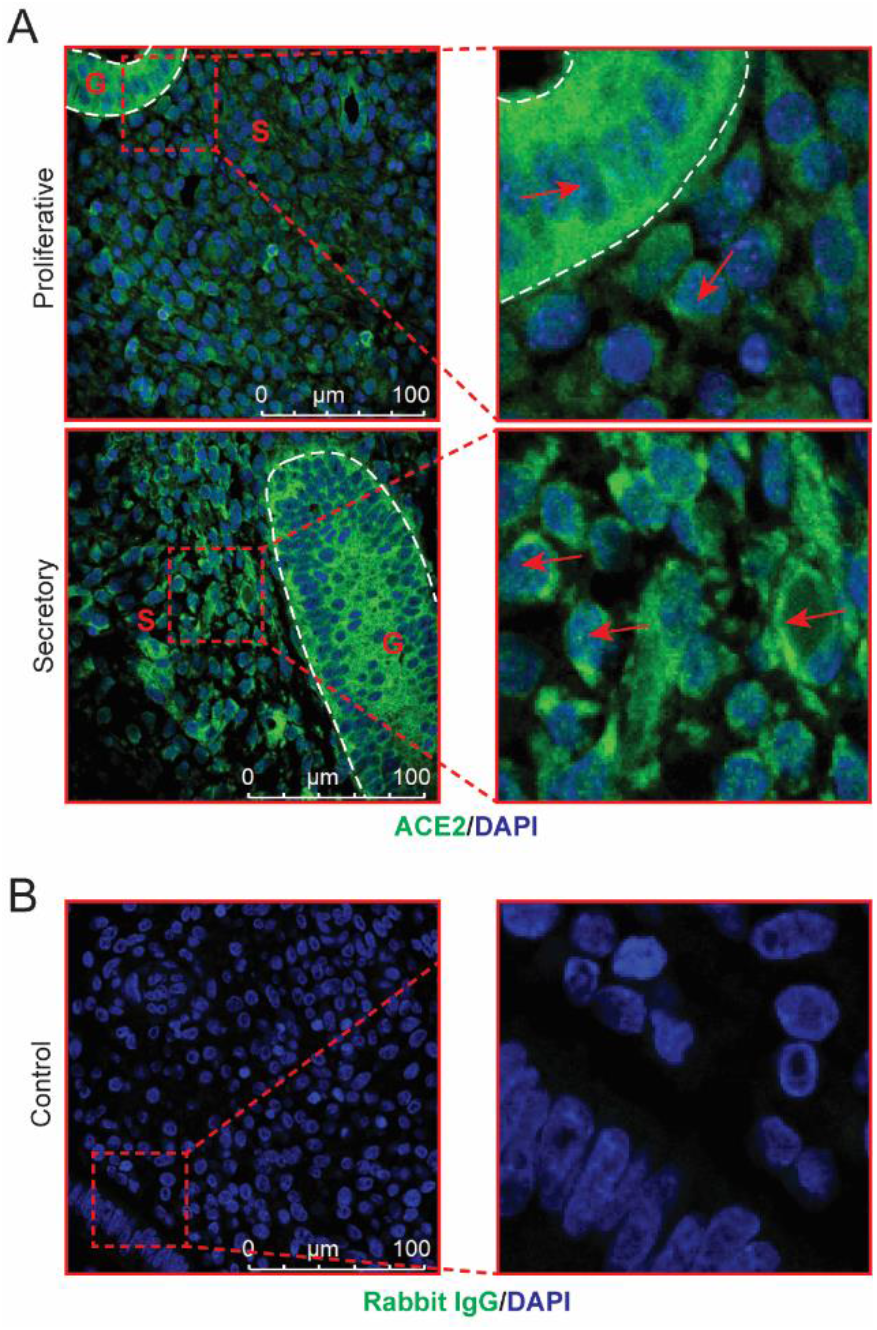
ACE2 protein expression is elevated in stromal cells of the secretory phase human endometrium. (**A**) Representative images showing immunolocalization of ACE2 (green) in proliferative (n=9) and secretory (n=6) phase endometrium. Blue stain is DAPI. G, gland; S, stroma. Red arrows indicate ACE2-positive cells, and the dashed white line marks the epithelium. (**B**) Rabbit IgG was used as an isotype control for staining. Scale bar: 100 μm.

**Figure 2:**
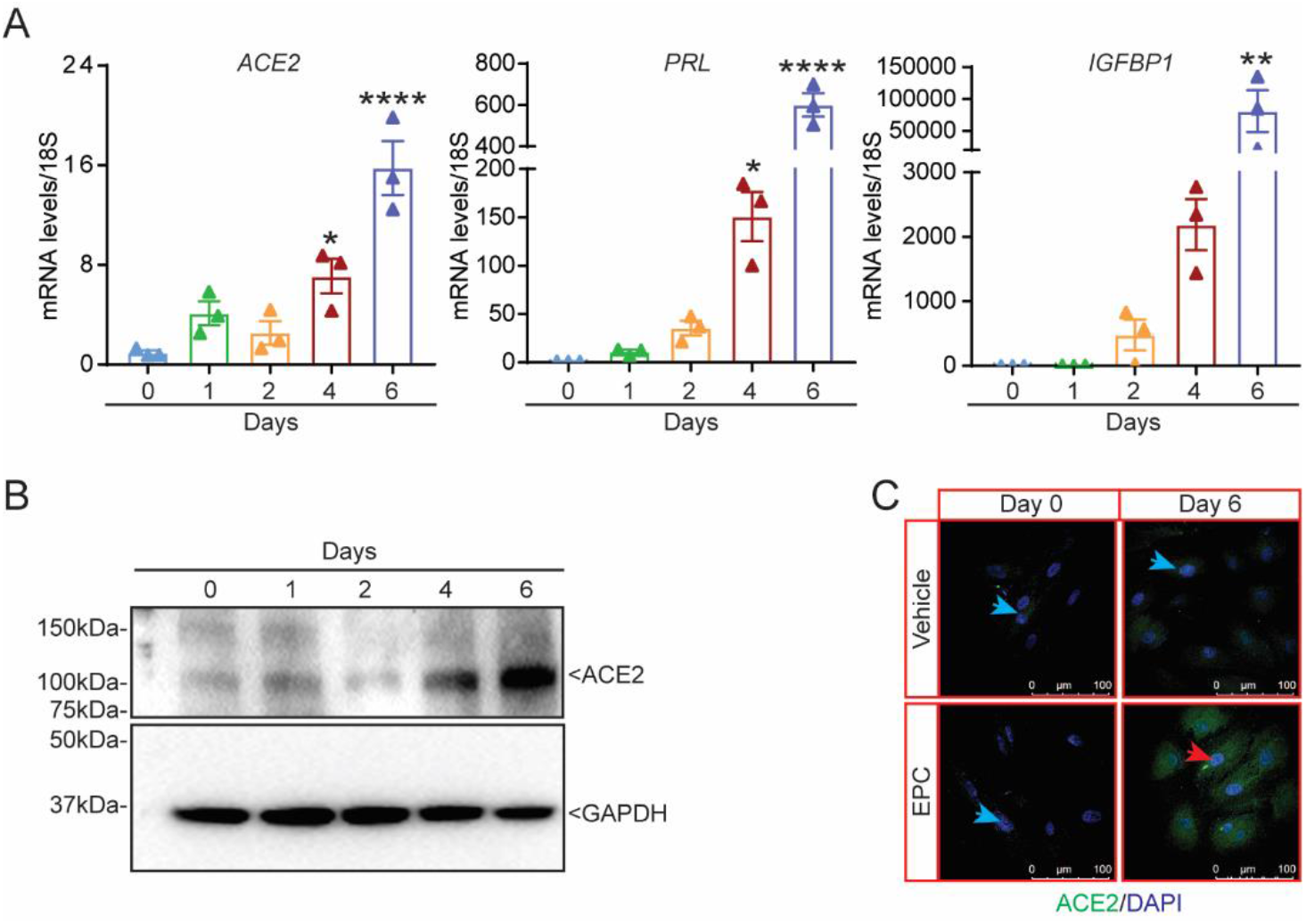
ACE2 is upregulated during *in vitro* human endometrial stromal cell decidualization. (**A**) Abundances of *ACE2, PRL*, and *IGFBP1* transcripts from human endometrial stromal cells (HESCs) induced to decidualize for the indicated numbers of days. Representative data from three replicates (n=3) from one subject sample are shown as mean ± SEM. The experiment were repeated three times. \**P* < 0.05, \*\**P* < 0.01, and \*\*\*\**P* < 0.0001. (**B**) Western blot of ACE2 from HESCs cultured in decidualization media for the indicated numbers of days; GAPDH was used as an internal loading control. (**C**) Immunofluorescence detection of ACE2 (green) in HESCs cultured with vehicle or decidualization media (EPC) for the indicated numbers of days. Blue stain is DAPI. Red arrowhead indicates a decidualized cell, and blue arrowheads indicate non-decidualized cells. Scale bar: 100 μm.

Next, we wondered whether ACE2 was required for primary HESC decidualization. To answer this question, we transfected HESCs with control or *ACE2*-targeting siRNAs and then exposed the cells to decidualization conditions. HESCs transfected with control siRNA changed from fibroblastic to epithelioid morphology (**Fig. 3A**) and had increased expression of the decidualization markers *PRL* and *IGFBP1* (**Fig. 3B**). In contrast, HESCs transfected with *ACE2*-targeting siRNA did not show a morphology change over six days (**Fig. 3A**) and expressed significantly less *PRL*, *IGFBP1*, and *ACE2* than control cells (**Fig. 3B-C**). These results demonstrate that ACE2 is essential for endometrial stromal cell decidualization.

**Figure 3:**
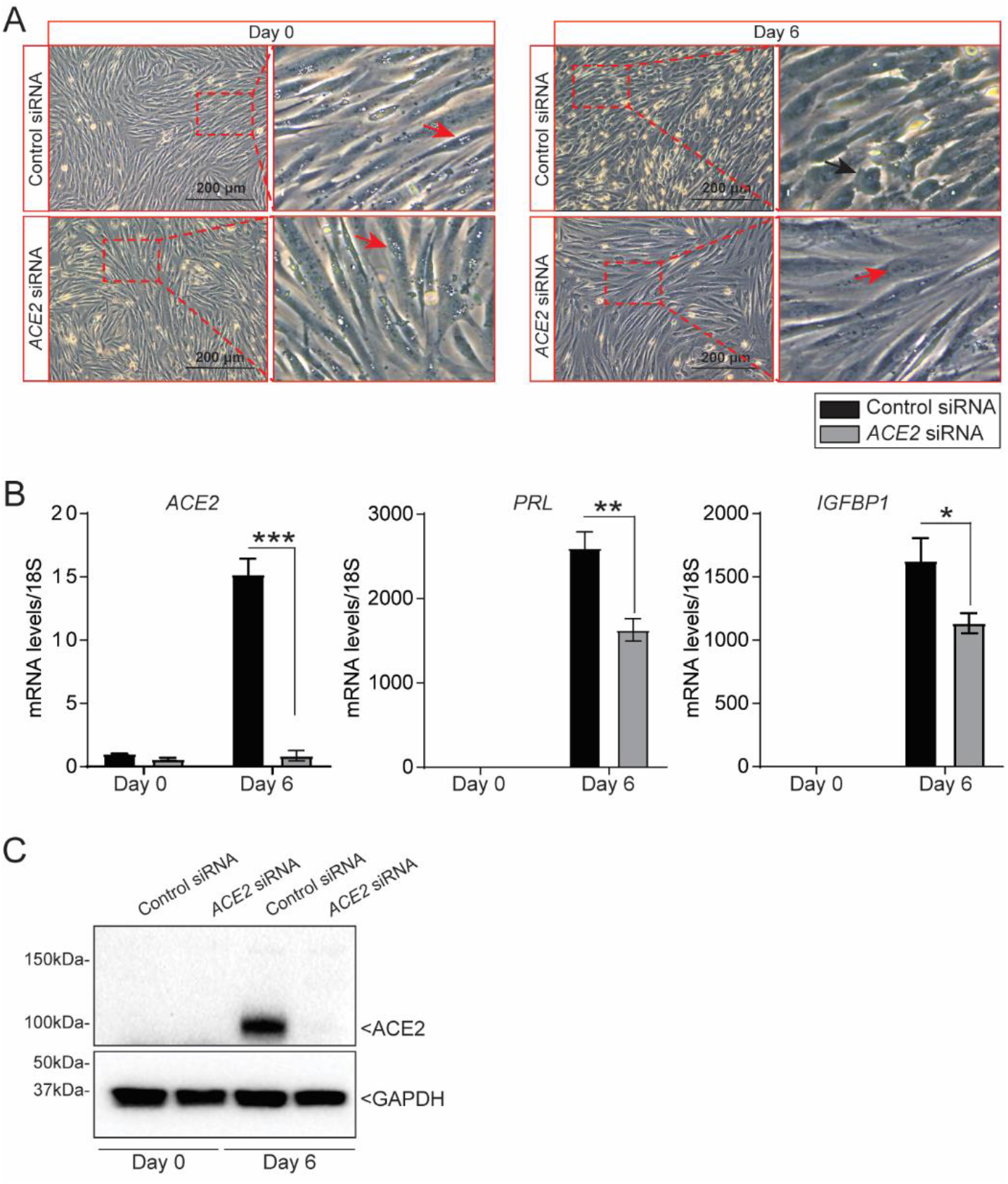
ACE2 is critical for human endometrial stromal cell decidualization. (**A**) Morphology of human endometrial stromal cells (HESCs) transfected with control or *ACE2* siRNA at day 0 or after six days of culture in decidualization conditions. Red arrows indicate non-decidualized cells, and the black arrow indicates a decidualized cell. Scale bar: 200 μm. (**B**) Abundances of *ACE2, PRL*, and *IGFBP1* transcripts in HESCs transfected with control or *ACE2* siRNAs and induced to decidualize for the indicated numbers of days. (**C**) Western blot of ACE2 protein from HESCs transfected with control or *ACE2* siRNA; GAPDH was used as an internal loading control. Representative data from three replicates from one subject sample are shown as mean ± SEM. The experiment was repeated three times; \**P* < 0.05, \*\**P* < 0.01, and \*\*\**P* < 0.001.

Finally, we examined the expression of ACE2 in the endometrium during early pregnancy in mice. We mated female wild-type mice with males of proven fertility and then stained their uteri with an ACE2-specific antibody at different days in early pregnancy. In days one through four, ACE2 localized to the cytoplasm and cell surface of epithelial and stromal cells. However, beginning on day three, strong ACE2 staining was seen in the cytoplasm of stromal cells. This staining was evident at least through day six, which is when robust decidualization occurs (**Fig. 4**). Given this change in ACE2 abundance during pregnancy, we wondered whether ACE2 expression was regulated by steroid hormones. To test this, we ovariectomized six-week-old mice, waited two weeks, treated the mice with either estrogen or progesterone for six hours, and then collected the uteri (**Fig. 5A**). Uteri from progesterone-treated mice expressed significantly more *Ace2* mRNA than uteri from vehicle-treated mice, which expressed significantly more *Ace2* mRNA than uteri from estrogen-treated (**Fig. 5B**). Consistent with this, immunofluorescence revealed that uteri from progesterone-treated mice had significantly more ACE2 protein in stromal cells than did uteri from vehicle- or estrogen-treated mice (**Fig. 5C**).

**Figure 4:**
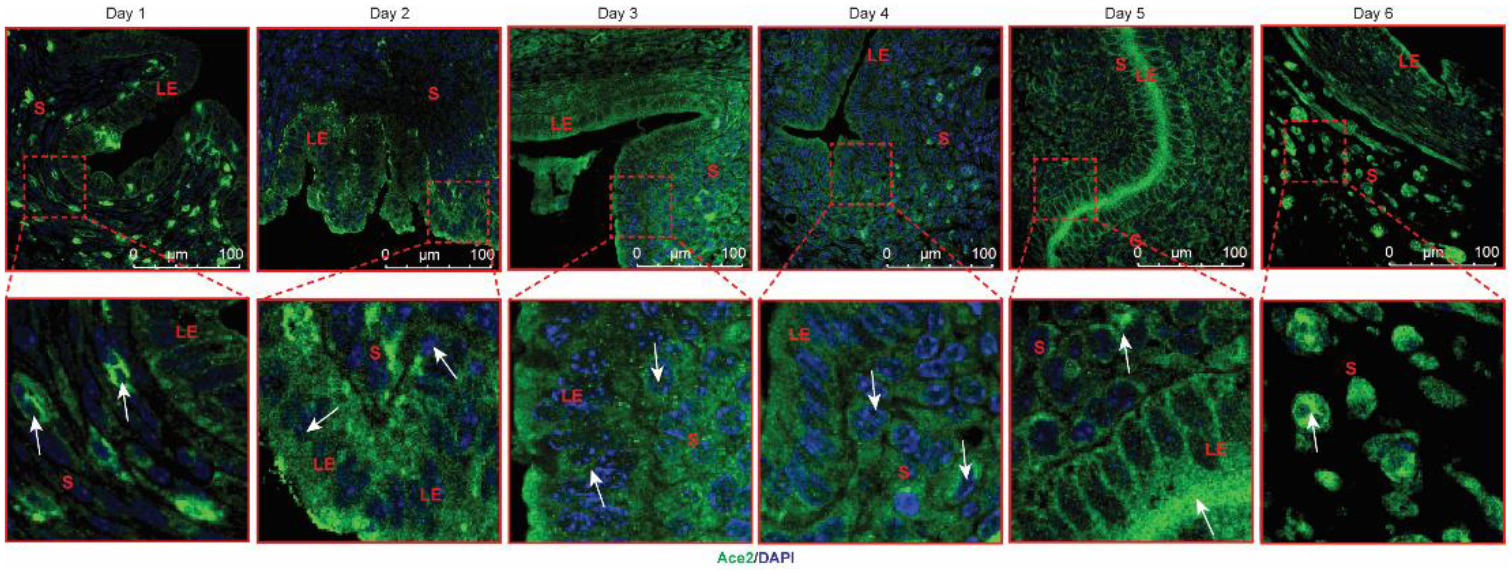
ACE2 protein expression in the mouse uterine stroma increases during early pregnancy. Shown are representative images of immunocytochemical localization of ACE2 in mouse uteri on the indicated days of pregnancy. LE, luminal epithelium; S, stroma; G, glands. Scale bar: 100 μm. White arrows indicate ACE2-positive cells. Samples from at least five mice were examined. All uteri were collected between 9:00 am and 10:00 am on the indicated days of pregnancy.

**Figure 5:**
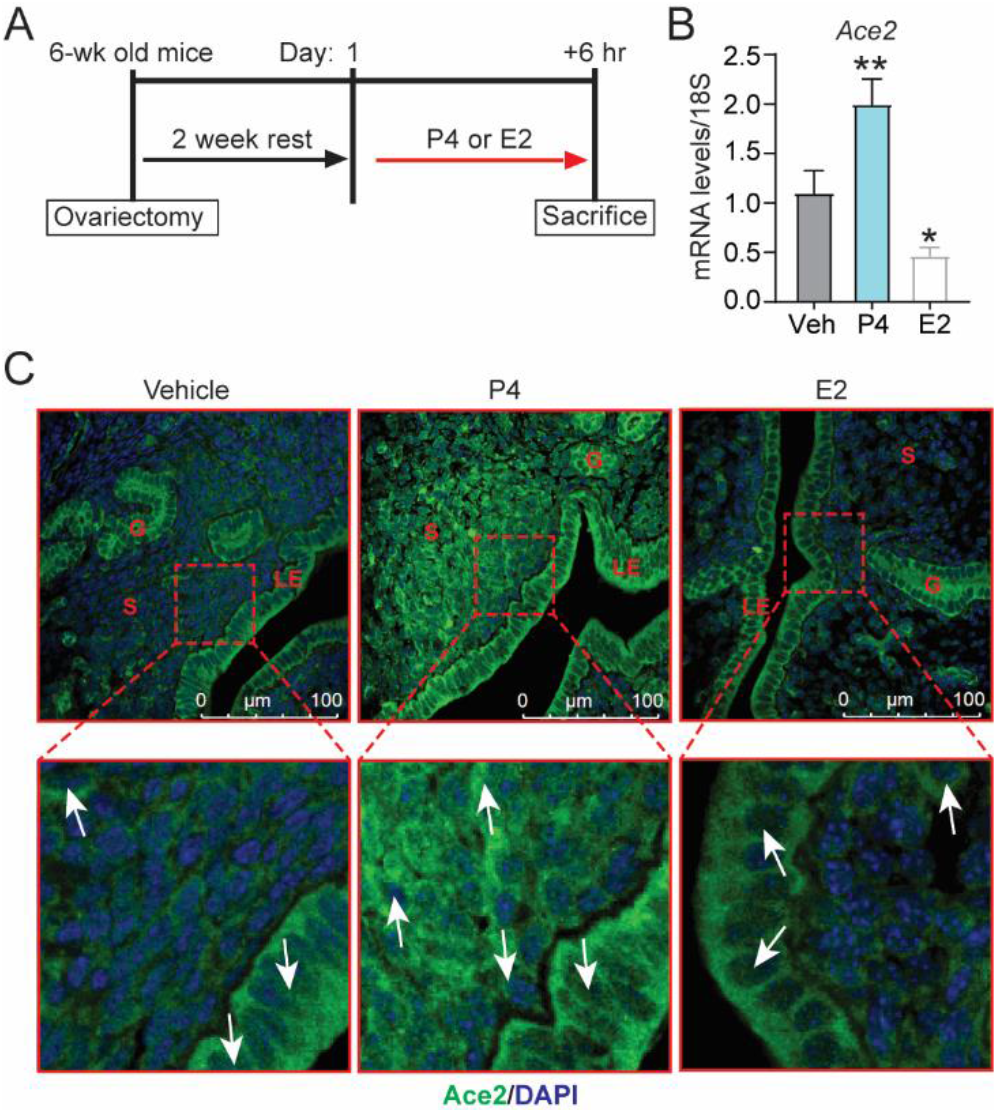
ACE2 expression in the mouse uterus is upregulated by progesterone exposure. (**A**) Experimental protocol and hormone treatment. E2, estrogen; P4, progesterone. Relative *Ace2* mRNA abundance after six hours of estrogen or progesterone treatment. Data are presented as mean ± SEM (n=5 mice per group). \**P*< 0.05, \*\**P*< 0.01. (**C**) Representative cross-sectional images of uteri stained for ACE2 (green) and DNA (blue); LE, luminal epithelium; S, stroma; G, glands, scale bar: 100 μm.

Together, our findings suggest that ACE2 expression in the endometrial stroma is promoted by progesterone in both humans and mice. Moreover, we show that ACE2 is required for human stromal cell decidualization. Given the high ACE2 expression in the human endometrium, SARS-CoV-2 may be able to enter endometrial stromal cells and elicit pathological manifestations in women with COVID-19. If so, women with COVID-19 may be at increased risk of early pregnancy loss. As more data become available, epidemiologists and obstetricians should focus on this important issue and determine whether women who intend to get pregnant should undergo additional health screenings during the COVID-19 pandemic.

## Materials and Methods

### Human ethical approval and endometrial stromal cell isolation

Informed consent was obtained in accordance with a protocol approved by the Washington University in St. Louis Institutional Review Board (IRB ID #: 201612127). Additionally, all work involving human subjects followed the guidelines of the World Medical Association Declaration of Helsinki. Human endometrial biopsies of healthy, reproductive-age women were collected during the proliferative phase (days 9 to 12) and secretory phase (days 14 to 26) of the menstrual cycle. HESCs were isolated as described previously (Camden et al. 2017; Michalski et al. 2018). Briefly, proliferative phase endometrial biopsies were minced with sterile scissors and then digested in DMEM/F12 medium containing 2.5 mg/ml collagenase (Sigma-Aldrich, Saint Louis, MO, USA) and 0.5 mg/ml DNase I (Sigma-Aldrich) for 1.5 hours at 37 °C. Then, detached cells were centrifuged at 800g for 2.5 min. collected, and layered over a Ficoll-Paque reagent layer and centrifuged for 30 min. at 400g (GE Healthcare Biosciences, Pittsburgh, PA) to remove lymphocytes. The HESC fraction from the top layer was collected and filtered through a 40 μm nylon cell strainer (BD Biosciences, Franklin Lakes, NJ). HESCs collected from the filtrate were suspended in DMEM/F-12 media containing 10% FBS, 100 U/ml penicillin, and 0.1 mg/ml streptomycin at 37 °C with 5% CO_2_. Independent HESC lines isolated from three patients were used for each experiment. Represented data are from a single patient with three technical replicates.

### Transfection and HESC decidualization

HESCs were grown in a six-well culture plate to 60%–70% confluence and transfected with 60 pmol of non-targeting siRNA (D-001810-10-05) or siRNAs targeting *ACE2* (L-005755-00-0005) (GE Healthcare Dharmacon Inc., Lafayette, CO) in Lipofectamine 2000 reagent (Invitrogen Corporation, Carlsbad, USA) as described previously (Camden et al. 2017). After 48 hours, HESCs were decidualized by culturing in EPC (Estrogen, Medroxy Progesterone Acetate and cAMP) medium (1x Opti-MEM reduced-serum media containing 2% FBS, 100 nM estradiol [cat. no. E1024, Sigma-Aldrich], 10 μM Medroxyprogesterone17-acetate [cat. no. M1629, Sigma-Aldrich], and 50 μM 8-Bromoadenosine 3′,5′-cyclic monophosphate sodium salt [cat. no. B7880, Sigma-Aldrich]). The EPC medium was changed every 48 hours until day six, when the cells were harvested for RNA isolation with the total RNA isolation kit (Invitrogen/Life Technologies, Grand Island, NY) or for protein isolation.

### Quantitative real-time PCR

Total RNA was extracted from uterine tissues or HESCs by using the total RNA isolation kit (Invitrogen/Life Technologies) according to the manufacturer's instructions. RNA was quantified with a Nano-Drop 2000 (Thermo Scientific, Waltham, MA, USA). Then, 1 μg of RNA was reverse transcribed with the High-Capacity cDNA Reverse Transcription Kit (Thermo Scientific, Waltham, MA, USA). The amplified cDNA was diluted to 10 ng/μl, and quantitative PCR was performed with primers specified in **Table S1** and Fast Taqman 2X mastermix (Applied Biosystems/Life Technologies, Grand Island, NY) on a 7500 Fast Real-time PCR system (Applied Biosystems/Life Technologies). Ribosomal RNA (18S) was used as an internal control for gene specific primers. (Camden et al. 2017; Kommagani et al. 2013; Kommagani et al. 2016).

### SDS-PAGE and Western blotting

Protein extracts were prepared from HESCs as described previously (Oestreich et al. 2020). Briefly, total proteins were extracted by homogenizing cells in RIPA lysis buffer (cat. no. 9806, Cell Signaling Technology) and centrifuging at 14,000 g for 15 minutes at 4 °C. The supernatants were collected and protein was quantified with the BCA Protein Assay kit according to the manufacturer’s instructions (Pierce BCA protein assay kit, cat no. 23227). Lysates containing 40 μg of protein were loaded on a 4-15% SDS-PAGE gel, separated with 1x Tris-Glycine Running Buffer, and transferred to PVDF membranes on a wet electro-blotting system (all from Bio-Rad, USA), all according to the manufacturer’s directions. The PVDF membranes were washed, blocked for 1 hour in 5% non-fat milk in TBS-T (Bio-Rad, USA), and incubated with primary antibodies anti-ACE2 (1:1000, ab15348, Abcam) and anti-GAPDH (1:3000, #2118S Cell Signaling Technology, USA) in 5% BSA in TBS-T overnight at 4°C. Then, blots were probed with anti-Rabbit IgG conjugated with horseradish peroxidase (1:5000, #7074, Cell Signaling Technology) in 5% BSA in TBS-T for 1 hour at room temperature. Signal was detected by using the Immobilon Western Chemiluminescent HRP Substrate (Millipore, MA, USA), and blot images were collected with a Bio-Rad ChemiDoc imaging system (Kommagani et al. 2016).

### Immunofluorescence

Formalin-fixed, paraffin-embedded sections (5 μm) of human endometrium and mouse uterus were deparaffinized in xylene, rehydrated in an ethanol gradient, and then boiled in antigen retrieval citrate buffer (Vector Laboratories Inc., CA, USA). Subsequently, sections were blocked with 2.5% goat serum in PBS (Vector laboratories) for one hour at room temperature, and then incubated overnight at 4°C with anti-ACE2 antibody (1:200, ab15348, Abcam) or normal rabbit IgG (#2729, Cell Signaling Technology). Then, sections were washed with PBS, incubated with Alexa Fluor 488-conjugated secondary antibody (Life Technologies) for one hour at room temperature, washed three times with PBS, and mounted with ProLong Gold Antifade Mountant with DAPI (cat. no. P36962 Thermo Scientific). Immunofluorescence images were captured on a confocal microscope (Leica DMI 4000B).

### Immunocytochemistry

HESCs were grown on poly-L-Lysine coated coverslips in 12-well plates and allowed to decidualize for six days in EPC media as described above. Then, cells were fixed with 4% paraformaldehyde (Alfa Aesar, USA) in PBS) for 20 min. at room temperature, washed with PBS, and permeabilized with 0.2% Triton X-100 (Sigma Aldrich, USA) in PBS for 20 min. at room temperature. Then, cells were washed, blocked with 2.5% normal goat-serum (Vector laboratories) in PBS for 1 h at room temperature, and incubated overnight at 4°C with anti-ACE2 antibody (ab15348, Abcam, 1:200) in 2.5% normal goat serum. Cells were washed and incubated with Alexa Fluor 488-conjugated secondary antibodies (Life Technologies) for one hour at room temperature and mounted with ProLong Gold Antifade Mountant with DAPI (Thermo Scientific). Images were captured on a confocal microscope (Leica DMI 4000B).

### Mice and hormone treatments

All mouse experimental procedures followed a protocol approved by the Washington University in St. Louis Institutional Animal Care and Use Committee (Protocol Number 20191079). CD1 wild-type mice (Charles River, Saint Louis, Missouri) were maintained on a 12-h light:12-h dark cycle. Sexually mature (8-week-old) CD1 females were mated to fertile wild-type males, and copulation was confirmed by the presence of vaginal plug on the following morning, designated as 1 day post-coital (dpc). Mice were euthanized, and uteri were collected on 1, 2, 3, 4, 5, and 6 dpc. To determine the uterine estrogen or progesterone responses, six-week-old CD1 mice were bilaterally ovariectomized, rested for two weeks to allow the endogenous ovarian-derived steroid hormones to dissipate, and then subcutaneously injected with 100 μl sesame oil (vehicle control), 1 mg progesterone, or 100 ng estradiol (Sigma-Aldrich) in 100 μl sesame oil. Six hours later, mice were euthanized, uterine tissues were collected and fixed in 4% paraformaldehyde, and RNA was isolated and processed for qRT-PCR (Kommagani et al. 2016).

### Statistical analyses

A two-tailed paired Student t-test was used to analyze experiments with two experimental groups, and analysis of variance by non-parametric alternatives was used for multiple comparisons to analyze experiments containing more than two groups. *P*<0.05 was considered significant. All data are presented as mean ± SEM. GraphPad Prism 8 software was used for all statistical analyses.

## Non-standard Abbreviations

SARS-CoV-2: severe acute respiratory syndrome coronavirus
ACE2: Angiotensin converting enzyme 2
WT: Wild Type
HESC: Human Endometrial Stromal Cells
dpc: Days Post Coitum
E2: Estrogen
P4: Progesterone

## Acknowledgments

We thank Dr. Deborah J. Frank (Department of Obstetrics and Gynecology, Washington University) for assistance with manuscript editing.

## Author contribution statement

RK conceived the project, supervised the work, analyzed the data, and wrote the manuscript. SBC, VKM, and PP, conducted the studies and wrote the manuscript. All authors reviewed and approved the final version of the manuscript.

## Funding

This work was funded, in part, by National Institutes of Health/National Institute of Child Health and Human Development grants R00HD080742 and RO1HD065435 to RK and Washington University School of Medicine start-up funds to RK.

## Declaration of Interest

The authors have no conflicts of interest to declare.

## Notes

The authors have declared that no conflict of interest exists relating to this work.

### Competing Interest Statement

The authors have declared no competing interest.

## References

Ali, M., et al. (2020), ‘The role of asymptomatic class, quarantine and isolation in the transmission of COVID-19’, J Biol Dyn, 14 (1), 389–408.

AlSaad, A. M. S., et al. (2020), ‘Renin angiotensin system blockage by losartan neutralize hypercholesterolemia-induced inflammatory and oxidative injuries’, Redox Rep, 25 (1), 51–58.

Baud, D., et al. (2020), ‘Second-Trimester Miscarriage in a Pregnant Woman With SARS-CoV-2 Infection’, JAMA.

Brosnihan, K. B., et al. (2012), ‘Decidualized pseudopregnant rat uterus shows marked reduction in Ang II and Ang-(1-7) levels’, Placenta, 33 (1), 17–23.

Camden, A. J., et al. (2017), ‘Growth regulation by estrogen in breast cancer 1 (GREB1) is a novel progesterone-responsive gene required for human endometrial stromal decidualization’, Mol Hum Reprod, 23 (9), 646–53.

Carson, D. D., et al. (2000), ‘Embryo implantation’, Dev Biol, 223 (2), 217–37.

Donoghue, M., et al. (2000), ‘A novel angiotensin-converting enzyme-related carboxypeptidase (ACE2) converts angiotensin I to angiotensin 1-9’, Circ Res, 87 (5), E1–9.

Favre, G., et al. (2020), ‘2019-nCoV epidemic: what about pregnancies?’, Lancet, 395 (10224), e40.

Fernandez, L. A., Twickler, J., and Mead, A. (1985), ‘Neovascularization produced by angiotensin II’, J Lab Clin Med, 105 (2), 141–5.

Gellersen, B. and Brosens, J. J. (2014), ‘Cyclic decidualization of the human endometrium in reproductive health and failure’, Endocr Rev, 35 (6), 851–905.

Goldenberg, I., et al. (2001), ‘Angiotensin II-induced apoptosis in rat cardiomyocyte culture: a possible role of AT1 and AT2 receptors’, J Hypertens, 19 (9), 1681–9.

Hamming, I., et al. (2004), ‘Tissue distribution of ACE2 protein, the functional receptor for SARS coronavirus. A first step in understanding SARS pathogenesis’, J Pathol, 203 (2), 631–7.

Harmer, D., et al. (2002), ‘Quantitative mRNA expression profiling of ACE 2, a novel homologue of angiotensin converting enzyme’, FEBS Lett, 532 (1-2), 107–10.

Jing, Y., et al. (2020), ‘Potential influence of COVID-19/ACE2 on the female reproductive system’, Mol Hum Reprod.

Khan, S., et al. (2020), ‘Emergence of a Novel Coronavirus, Severe Acute Respiratory Syndrome Coronavirus 2: Biology and Therapeutic Options’, J Clin Microbiol, 58 (5).

Kommagani, R., et al. (2016), ‘The Promyelocytic Leukemia Zinc Finger Transcription Factor Is Critical for Human Endometrial Stromal Cell Decidualization’, PLoS Genet, 12 (4), e1005937.

Kommagani, R., et al. (2013), ‘Acceleration of the glycolytic flux by steroid receptor coactivator-2 is essential for endometrial decidualization’, PLoS Genet, 9 (10), e1003900.

Le Noble, F. A., et al. (1993), ‘Evidence for a novel angiotensin II receptor involved in angiogenesis in chick embryo chorioallantoic membrane’, Am J Physiol, 264 (2 Pt 2), R460–5.

Merrill, D. C., et al. (2002), ‘Angiotensin-(1-7) in normal and preeclamptic pregnancy’, Endocrine, 18 (3), 239–45.

Michalski, S. A., et al. (2018), ‘Isolation of Human Endometrial Stromal Cells for In Vitro Decidualization’, J Vis Exp, (139).

Muus, Christoph, et al. (2020), ‘Integrated analyses of single-cell atlases reveal age, gender, and smoking status associations with cell type-specific expression of mediators of SARS-CoV-2 viral entry and highlights inflammatory programs in putative target cells’, bioRxiv, 2020.04.19.049254.

Nakashima, H., et al. (2006), ‘Angiotensin II regulates vascular and endothelial dysfunction: recent topics of Angiotensin II type-1 receptor signaling in the vasculature’, Curr Vasc Pharmacol, 4 (1), 67–78.

Neves, L. A., et al. (2008), ‘ACE2 and ANG-(1-7) in the rat uterus during early and late gestation’, Am J Physiol Regul Integr Comp Physiol, 294 (1), R151–61.

Norwitz, E. R., Schust, D. J., and Fisher, S. J. (2001), ‘Implantation and the survival of early pregnancy’, N Engl J Med, 345 (19), 1400–8.

Oestreich, A. K., et al. (2020), ‘The Autophagy Gene Atg16L1 is Necessary for Endometrial Decidualization’, Endocrinology, 161 (1).

Okada, H., Tsuzuki, T., and Murata, H. (2018), ‘Decidualization of the human endometrium’, Reprod Med Biol, 17 (3), 220–27.

Okada, H., et al. (2014), ‘Regulation of decidualization and angiogenesis in the human endometrium: mini review’, J Obstet Gynaecol Res, 40 (5), 1180–7.

Peach, M. J. (1977), ‘Renin-angiotensin system: biochemistry and mechanisms of action’, Physiol Rev, 57 (2), 313–70.

Qin, S., et al. (2013), ‘[Expression and significance of ACE2-Ang-(1-7)-Mas axis in the endometrium of patients with polycystic ovary syndrome]’, Zhonghua Yi Xue Za Zhi, 93 (25), 1989–92.

Riviere, G., et al. (2005), ‘Angiotensin-converting enzyme 2 (ACE2) and ACE activities display tissue-specific sensitivity to undernutrition-programmed hypertension in the adult rat’, Hypertension, 46 (5), 1169–74.

Santos, R. A., et al. (2003), ‘Angiotensin-(1-7) is an endogenous ligand for the G protein-coupled receptor Mas’, Proc Natl Acad Sci U S A, 100 (14), 8258–63.

Schaller, T., et al. (2020), ‘Postmortem Examination of Patients With COVID-19’, JAMA.

Schwartz, D. A. and Dhaliwal, A. (2020), ‘INFECTIONS IN PREGNANCY WITH COVID-19 AND OTHER RESPIRATORY RNA VIRUS DISEASES ARE RARELY, IF EVER, TRANSMITTED TO THE FETUS: EXPERIENCES WITH CORONAVIRUSES, HPIV, hMPV RSV, AND INFLUENZA’, Arch Pathol Lab Med.

Shanes, E. D., et al. (2020a), ‘Placental pathology in COVID-19’, medRxiv.

Shanes, E. D., et al. (2020b), ‘Placental Pathology in COVID-19’, Am J Clin Pathol, 154 (1), 23–32.

Shang, J., et al. (2020), ‘Cell entry mechanisms of SARS-CoV-2’, Proc Natl Acad Sci U S A, 117 (21), 11727–34.

Tipnis, S. R., et al. (2000), ‘A human homolog of angiotensin-converting enzyme. Cloning and functional expression as a captopril-insensitive carboxypeptidase’, J Biol Chem, 275 (43), 33238–43.

Vaz-Silva, J., et al. (2009), ‘The vasoactive peptide angiotensin-(1-7), its receptor Mas and the angiotensin-converting enzyme type 2 are expressed in the human endometrium’, Reprod Sci, 16 (3), 247–56.

Verdecchia, P., et al. (2020), ‘The pivotal link between ACE2 deficiency and SARS-CoV-2 infection’, Eur J Intern Med, 76, 14–20.

Vickers, C., et al. (2002), ‘Hydrolysis of biological peptides by human angiotensin-converting enzyme-related carboxypeptidase’, J Biol Chem, 277 (17), 14838–43.

Wang, H. and Dey, S. K. (2006), ‘Roadmap to embryo implantation: clues from mouse models’, Nat Rev Genet, 7 (3), 185–99.

Wilcox, A. J., Baird, D. D., and Weinberg, C. R. (1999), ‘Time of implantation of the conceptus and loss of pregnancy’, N Engl J Med, 340 (23), 1796–9.

Wrapp, D., et al. (2020), ‘Cryo-EM structure of the 2019-nCoV spike in the prefusion conformation’, Science, 367 (6483), 1260–63.

Yamada, T., Horiuchi, M., and Dzau, V. J. (1996), ‘Angiotensin II type 2 receptor mediates programmed cell death’, Proc Natl Acad Sci U S A, 93 (1), 156–60.

Yuan, J., et al. (2018), ‘Tridimensional visualization reveals direct communication between the embryo and glands critical for implantation’, Nat Commun, 9 (1), 603.

Yuan, J., et al. (2019), ‘Primary decidual zone formation requires Scribble for pregnancy success in mice’, Nat Commun, 10 (1), 5425.

Zhang, C., et al. (2020), ‘Impact of population movement on the spread of 2019-nCoV in China’, Emerg Microbes Infect, 9 (1), 988–90.

Zhou, P., et al. (2020), ‘A pneumonia outbreak associated with a new coronavirus of probable bat origin’, Nature, 579 (7798), 270–73.

