## Supplemental Table 1 for "The SARS-CoV-2 receptor, Angiotensin converting enzyme 2 (ACE2) is required for human endometrial stromal cell decidualization"

Table S1. List of TaqMan probes

| Gene name | Species | Application, Chemistry | Company | Cat. No. |
|---|---|---|---|---|
| *PRL* | Human | qPCR, Taqman | ABI | Hs00168730_m1 |
| *IGFBP1* | Human | qPCR, Taqman | ABI | Hs00236877_m1 |
| *ACE2* | Human | qPCR, Taqman | ABI | Hs01085333_m1 |
| *Ace2* | Mouse | qPCR, Taqman | ABI | Mm01159006_m1 |
| *18S* | Mouse | qPCR, Taqman | ABI | 4318839 |

ABI-applied biosystems
